# Angiotensin-II drives changes in microglia-vascular interactions in rats with heart failure

**DOI:** 10.1101/2023.12.22.573045

**Authors:** Ferdinand Althammer, Ranjan K. Roy, Matthew K. Kirchner, Shaina McGrath, Elba Campos Lira, Javier E. Stern

## Abstract

Activation of microglia, the resident immune cells of the central nervous system, leading to the subsequent release of pro-inflammatory cytokines, has been linked to cardiac remodeling, autonomic disbalance, and cognitive deficits in heart failure (HF). While previous studies emphasized the role of hippocampal Angiotensin II (AngII) signaling in HF-induced microglial activation, unanswered mechanistic questions persist. Evidence suggests significant interactions between microglia and local microvasculature, potentially affecting blood-brain barrier integrity and cerebral blood flow regulation. Still, whether the microglial-vascular interface is affected in the brain during HF remains unknow. Using a well-established ischemic HF rat model, we demonstrate increased vessel-associated microglia (VAM) in HF rat hippocampi, which showed heightened expression of AngII AT1a receptors. Acute AngII administration to sham rats induced microglia recruitment to the perivascular space, along with increased expression of TNFa. Conversely, administering an AT1aR blocker to HF rats prevented the recruitment of microglia to the perivascular space, normalizing their levels to those in healthy rats. These results highlight the critical importance of a rather understudied phenomenon (i.e., microglia-vascular interactions in the brain) in the context of the pathophysiology of a highly prevalent cardiovascular disease, and unveil novel potential therapeutic avenues aimed at mitigating neuroinflammation in cardiovascular diseases.

## Introduction

Heart failure (HF) is a complex clinical syndrome characterized by the inability of the heart to adequately pump blood and meet the body’s metabolic demands^1^. It represents a significant public health concern worldwide due to its high prevalence, morbidity, mortality, and associated healthcare costs ^1,2^. The burden of HF on society is substantial. According to the World Health Organization (WHO), an estimated 65 million people are affected by HF globally, and its prevalence is projected to increase due to aging populations and the rising prevalence of risk factors like obesity and diabetes ^3,4^. Despite the clinical and economic impact of HF on society ^2,3,5^, precise mechanisms explaining the manifold peripheral, central and systemic effects of this disease, as well as novel (sub-)cellular therapeutic targets are missing.

Research has increasingly highlighted the association between HF and cognitive deficits ^6–9^, with a spotlight on the hippocampus ^6,10–13^, a key player in memory and cognitive functions ^14^. Studies have demonstrated that HF patients often exhibit structural and functional alterations in the hippocampus ^6,12^, contributing to cognitive impairment. The hippocampus is particularly vulnerable due to its sensitivity to changes in cerebral blood flow ^15,16^ and its susceptibility to the effects of neuroinflammation ^17,18^. Emerging evidence suggests that neuroinflammation and microglial activation play a significant role in the progression of various cardiovascular diseases, including HF ^19–24^. Microglia, the resident immune cells of the CNS, are known to become activated in response to injury, infection, or inflammation ^25–27^. Importantly, their activation and the consequent release of pro-inflammatory cytokines has been shown to contribute to the progression of HF by affecting autonomic control, cardiac remodeling, and neuronal dysfunction ^19,23,24^. The early stage of HF involves a myriad of different signaling cascades. One of the most important and best-studied is the Angiotensin II (AngII)-AT1a receptor (AT1aR) cascade^1,19,23,28–32^, which becomes dysregulated, leading to excessive AngII production and elevated circulating AngII levels ^1,28,32–37^.

Notably, investigations into the role of microglial AngII signaling ^19,38–40^ have unveiled a potential mechanism underlying these cognitive deficits. Microglia, activated by the elevated levels of AngII in HF, can induce neuroinflammation in the hippocampus, disrupting synaptic plasticity and impairing memory consolidation ^10,19^. These findings emphasize the intricate interplay between cardiovascular health and cognitive function, with the hippocampus serving as a focal point for understanding and addressing cognitive deficits in HF patients. We recently demonstrated that microglial AngII-AT1aR signaling precedes cytokine production in the hippocampus of HF rats, and that administration of the AT1aR antagonist losartan improved various cellular, molecular, and behavioral endpoints in HF rats ^19^. This interaction between AngII-AT1aR signaling, neuroinflammation, and cardiac dysfunction underscores the intricate link between the cardiovascular and central nervous systems in HF.

Microglia are not only involved in neuroinflammation but also play a crucial role interacting with the local microvasculature. For example, microglia play a critical role in maintaining the integrity of the blood-brain barrier (BBB) ^25,41–45^, a selectively permeable barrier that separates the CNS from the systemic circulation, regulating the exchange of molecules between the two compartments ^42–44,46^. Moreover, a growing body of evidence supports a direct role of microglia in regulating brain blood flow ^47,48^. In a recent study, Bisht and colleagues used *in vivo* 2-photon imaging to monitor microglial movements in the mouse cortex and found that microglia-vessel interactions are highly dynamic ^48^. Moreover, Haruwaka and colleagues showed that during systemic inflammation, microglia migrate towards blood vessels and promote BBB stability through Claudin-5 ^41^. However, upon sustained inflammation, microglia begin to phagocytose astrocytic AQP4-positive endfeet in a CD68-dependent manner, thereby compromising BBB integrity ^41^. Thus, these studies support the notion that microglia-vascular interactions are highly dynamic, both under physiological and pathological conditions. Still, whether the microglial-vascular interface in the hippocampus is affected in HF, and whether the AngII-AT1aR signaling cascade is a major contributing factor, remains completely unknown.

To address this major gap in our knowledge, in the current study we used a well-established rat model of ischemic HF and employed a multidisciplinary approach including immunohistochemistry, RNAScope hybridization, three-dimensional microglia/vascular reconstruction, *in vivo* intracarotid infusion of vessel markers and fluorescently-labeled AngII, and administration of the AT1aR antagonist losartan. We specifically tested the hypothesis that the AngII-AT1aR signaling cascade contributes to dysregulation of the microglial-vascular interface in the hippocampus of rats with HF.

## Results

### Increased number of vessel-associated microglia (VAM) in HF rats

To explore the role of microglia-vessel interactions^41,48^ and to determine whether these interactions were altered in HF, we first stained brain sections containing the dorsal hippocampus (DH) of sham and HF rats with antibodies against the microglial marker IBA1 and the vascular marker aquaporin-4 (AQP4^41^) to investigate potential differences in the number of parenchymal vs. vessel-associated microglia (VAM) between the two groups (**Figure 1a, b**, complete echocardiographic assessment of sham and HF rats can be found in **Table 1**). We observed a significant increase in the number of VAM in HF rats (2.1-fold, **Figure 1c**), which was accompanied by an expected reduction of parenchymal microglia (**Figure 1d**). When we analyzed the morphological properties of VAM in detail using Imaris algorithms (see Methods), we observed considerable variation among cells reflected by alterations in overall shape and density of microglia-vessel contacts. Based on these quantitative properties, we further classified VAM into three subgroups (**Figure 1e**): Type I (rounded soma with only microglial filaments in contact with the vessel), Type II (rounded soma with both somatic and filament contact with the vessel) and Type III (elongated soma entirely in contact with the associated vessel). Intriguingly, we found that the relative abundance of each individual VAM subtype was essentially reversed in HF rats. While Type I VAM accounted for 65.2% and Type III for 10.5% in sham rats, HF rats displayed only 15.0% Type I, but 58.6% Type III microglia (Chi-square test, p<0.0001, **Figure 1f**).

**Figure 1.**
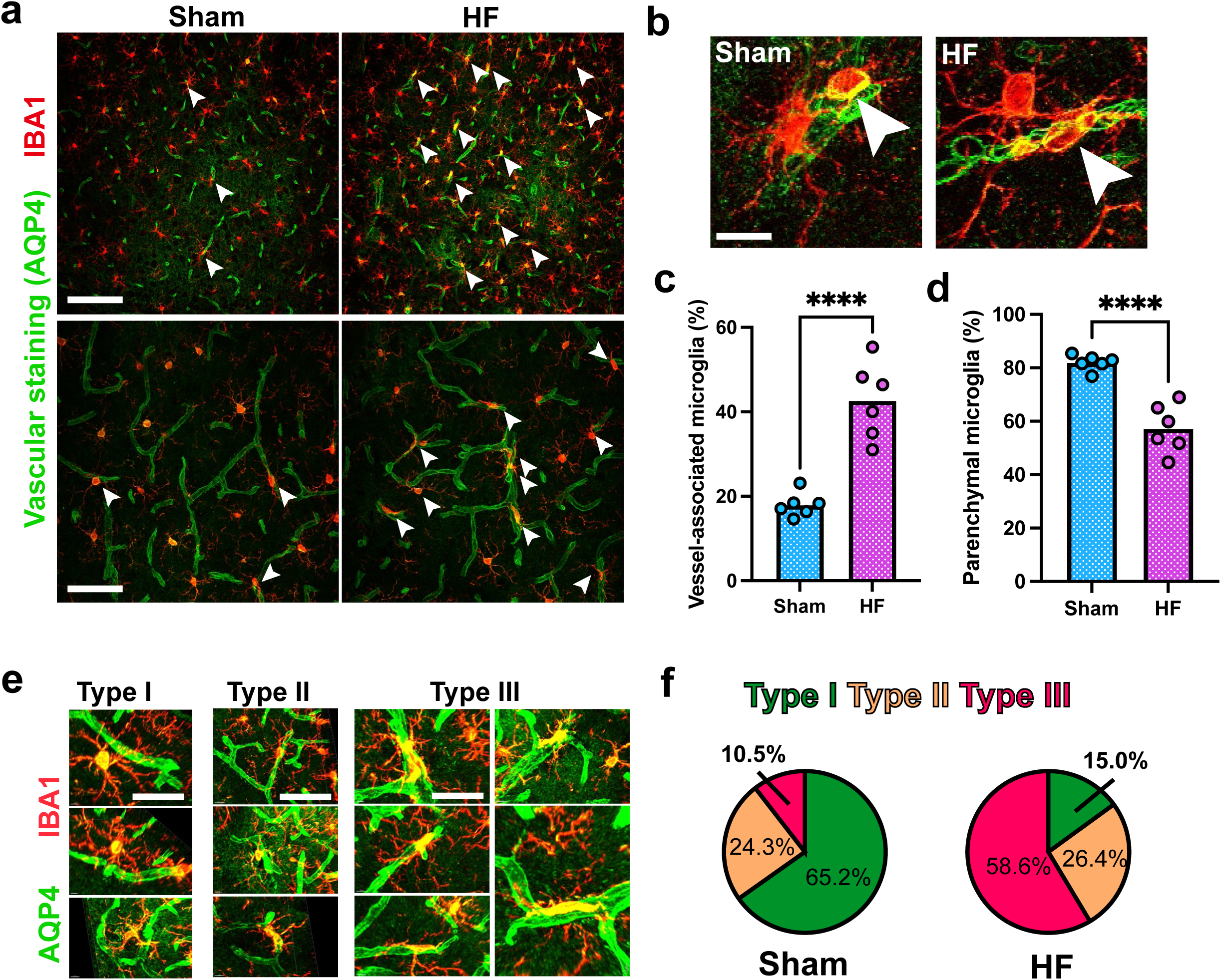
Increase in the number of vessel-associated microglia in HF rats. **a** Confocal images show the distribution of parenchymal and vessel-associated microglia in the dorsal hippocampus of sham and HF rats. Immunohistochemical staining against IBA1 (red) and the vessel marker AQP4 (green); white arrowheads indicate vessel-associated microglia. Scale bars 200µm, 50µm. **b** High magnification confocal images highlight vessel-associated microglia in sham and HF rats. Scale bar 10µm. **c** Quantification of vessel-associated and **d** parenchymal microglia in sham and HF rats (n=6 rats per group). ****p<0.0001 **e** Exemplary confocal images of Type I-III vessel-associated microglia, which are characterized based on the intensity of microglia-vessel contacts. Type I microglia contact blood vessels exclusively with their filaments, Type II microglia display somatic contact with the blood vessel and Type III seem to merge with the blood vessel along the entirety of the microglial soma. Scale bars, Type I: 25µm, Type II: 50µm, Type II: 25µm. **f** Pie charts highlight the respective percentage of Type I-III vessel-associated microglia (sham/HF pooled).

**Table 1.** Echocardiographic assessment of sham and HF rats used in this study.

|  | Sham | HF | p value |
| --- | --- | --- | --- |
| Heart Rate (BPM) | $297.53 \pm 31.19$ | $270.88 \pm 24.44$ | n.s. |
| Stroke Volume (uL) | $239.28 \pm 34.89$ | $126.03 \pm 15.74$ | **** |
| Ejection Fraction (%) | $82.27 \pm 3.2$ | $25.22 \pm 6.29$ | **** |
| Fractional Shortening (%) | $50.68 \pm 3.92$ | $14.36 \pm 1.28$ | **** |
| Cardiac Output (mL/min) | $70.85 \pm 10.82$ | $34.23 \pm 5.73$ | **** |
| LV Mass (mg) | $1846.48 \pm 105.62$ | $966.5 \pm 113.42$ | **** |
| Lvaw,d (mm) | $3.04 \pm 0.54$ | $1.46 \pm 0.29$ | **** |
| Lvaw,s (mm) | $3.67 \pm 0.40$ | $15.61 \pm 56.36$ | **** |
| Lvid,d (mm) | $6.9 \pm 0.57$ | $12.3 \pm 1.37$ | **** |
| Lvid,s (mm) | $4.43 \pm 0.54$ | $8.23 \pm 0.69$ | **** |
| Lvpw,d (mm) | 3.58±0.47 | 1.55±0.35 | **** |
| Lvpw,s (mm) | 3.68±0.26 | 1.93±0.26 | **** |

In addition, following a three-dimensional reconstruction of microglia and blood vessels, we observed frequent penetration of microglial filaments into the vessel lumen (**Figure 2a**). The frequency of vessel-protruding microglia (i.e. microglia with at least one protruding filament) was significantly increased in HF rats (3-fold, **Figure 2b**). Interestingly, we found that VAM that extended their processes into the vessel lumen belonged almost exclusively to Type III, which was consistent for both sham and HF rats (**Figure 2c**). Of note, a similar observation of vessel penetration by microglial processes has recently been reported by another group in the cerebral cortex of mice under baseline conditions^41^.

**Figure 2.**
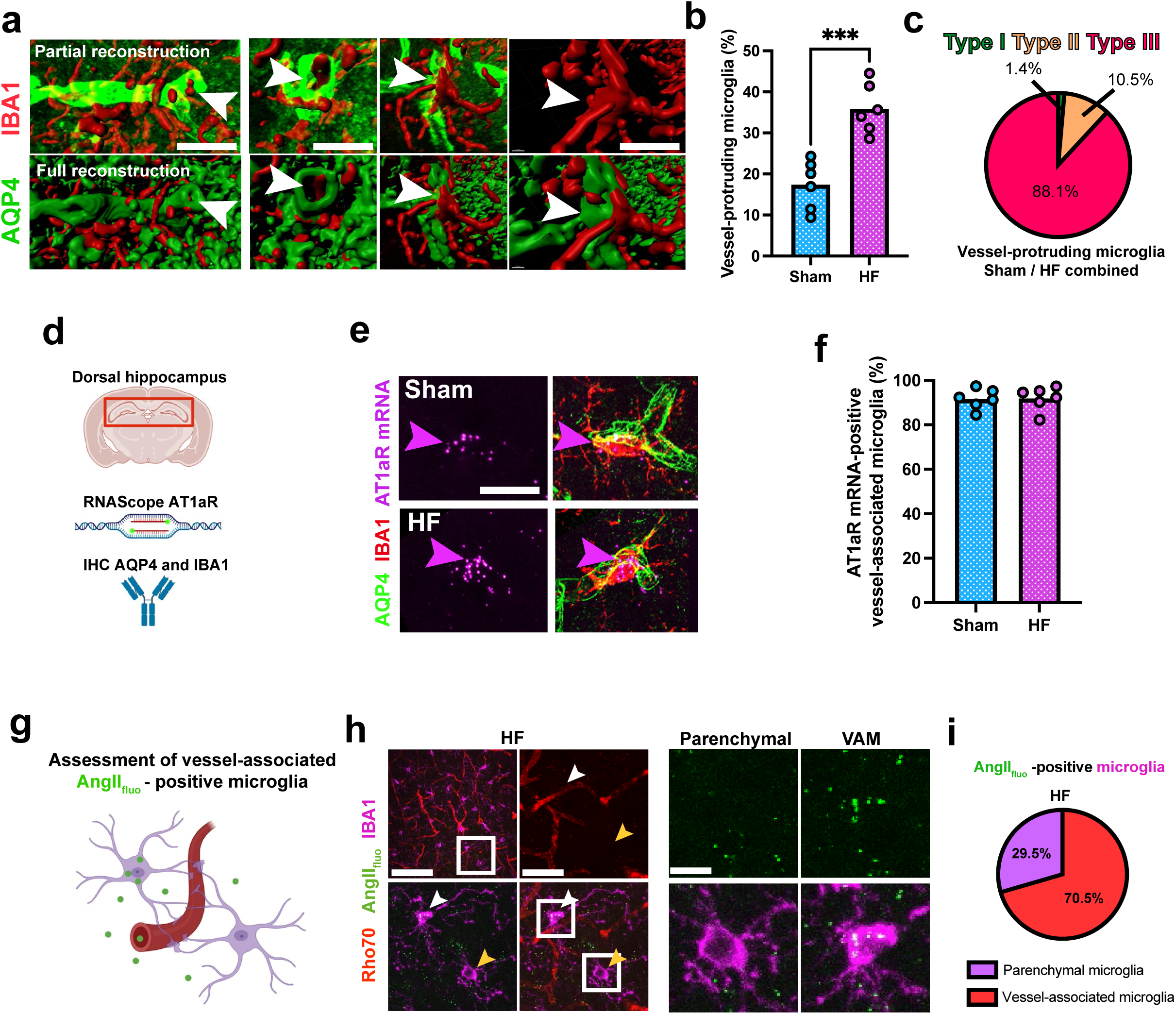
Vessel-protruding microglia are AT1aR mRNA-positive in both sham and HF rats. **a** Confocal images after partial and full three-dimensional reconstruction highlighting vessel protrusion by microglial filaments. White arrowheads indicate multiple instances of vessel-penetrating microglial filaments or soma. Scale bars 20µm, 10µm and 5µm. **b** Quantification of vessel-protruding microglia in sham and HF rats (n=6 rats per group). **c** Pie chart shows that vessel-protruding microglia almost exclusively occurs by Type III microglia. **d** Schematic illustration of the methodological approach to label and identify AT1aR mRNA-positive, vessel-associated microglia in the dorsal hippocampus of sham and HF rats. **e** Confocal images of AT1aR-positive, vessel-associated microglia in sham and HF rats; magenta arrowheads highlight co-localization of IBA1 and AT1aR mRNA. Scale bar 10µm. **f** Quantification of AT1aR-positive, vessel-associated microglia in sham and HF rats. **g** Schematic depiction of intracarotid infusions and AngII_fluo_ uptake by microglia. Confocal images of IBA-stained, AngII_fluo_-infused microglia were used for three-dimensional reconstruction and analysis. **h** Confocal images of a typical hippocampus of a HF infused with Rho70 and AngII_fluo_ stained for IBA1. Enlarged insets show a representative example of a VAM and parenchymal microglia, white arrowhead indicates VAM, yellow arrowhead indicates a parenchymal microglia. Enlarged insets (right) show co-localization of AngII_fluo_ in VAM but not parenchymal microglia. Scale bar 100µm, 25µm and 5µm. **i** Pie charts summarize the percentage of parenchymal and vessel-associated microglia, AngII_fluo_-positive microglia in HF rats.

### AT1aRs overexpression and leaked AngII levels in HF rats are restricted to VAM

Circulating AngII levels dramatically increase in HF^1,32^. Moreover, we recently showed that AngII can gain access to the hippocampal parenchyma in HF rats^19^, which is then taken up by microglia that overexpress AngII AT1a receptors (AT1aRs), initiating a neuroinflammatory cascade leading to neuronal deterioration and apoptosis in HF rats^19^. However, it remained unclear whether AT1R overexpression and leaked AngII uptake during HF occurs in parenchymal microglia or VAM. To this end, we first combined immunohistochemical staining to label microglia and vessels with RNAScope for AT1aR mRNA (**Figure 2d,e**). Intriguingly, our results show that virtually all VAM were AT1aR mRNA-positive, both in sham and HF rats (**Figure 2f**), indicating that the increased expression of AT1aRs in HF rats is mostly confined to this population of microglial cells. Based on this finding, we then hypothesized that the leaked AngII molecules in HF would be predominantly confined to the AT1aR-expressing VAM. To this end, we performed a new set of experiments in which we performed *in vivo* intracarotid infusions fluorescently-labeled AngII (AngII_fluo_) combined with the fluorescent dye Rhodamine70kDa (Rho70) to label the DH microvasculature, and assessed the location of the leaked AngII_flu_ in microglial subtypes. As expected, we found the majority of the leaked AngII_fluo_ in HF rats to be predominantly located in VAM (**Figure 2g-i**).

### AngII acting on AT1aRs is a critical signal contributing to recruitment of microglial to the perivascular space

To determine whether AngII is a key signal that could trigger the recruitment of parenchymal microglia into VAM, we performed a two-pronged approach. In the first set of experiments, we assessed the effects of an acute increase in circulating AngII levels. To this end, we performed *in vivo* intracarotid infusions of AngII (or saline as control) combined with the fluorescent dye Rhodamine70kDa (Rho70) to label the DH microvasculature and allowed both substances to circulate for 30 mins (**Figure 3a**). Processed brain sections of sham and HF rats were then used for immunohistochemical staining to quantify the number of parenchymal microglia and VAM (**Figure 3b**). Unexpectedly, we observed that the acute infusion of AngII significantly increased the number of VAM in sham rats (1.8-fold). In HF rats, as we reported above, the number of VAM was already significantly (2.6-fold) higher compared to sham rats. Still, AngII further increased the proportion of VAM, albeit to a lesser extent than sham rats (HF: 1.3-fold, **Figure 3c**).

**Figure 3.**
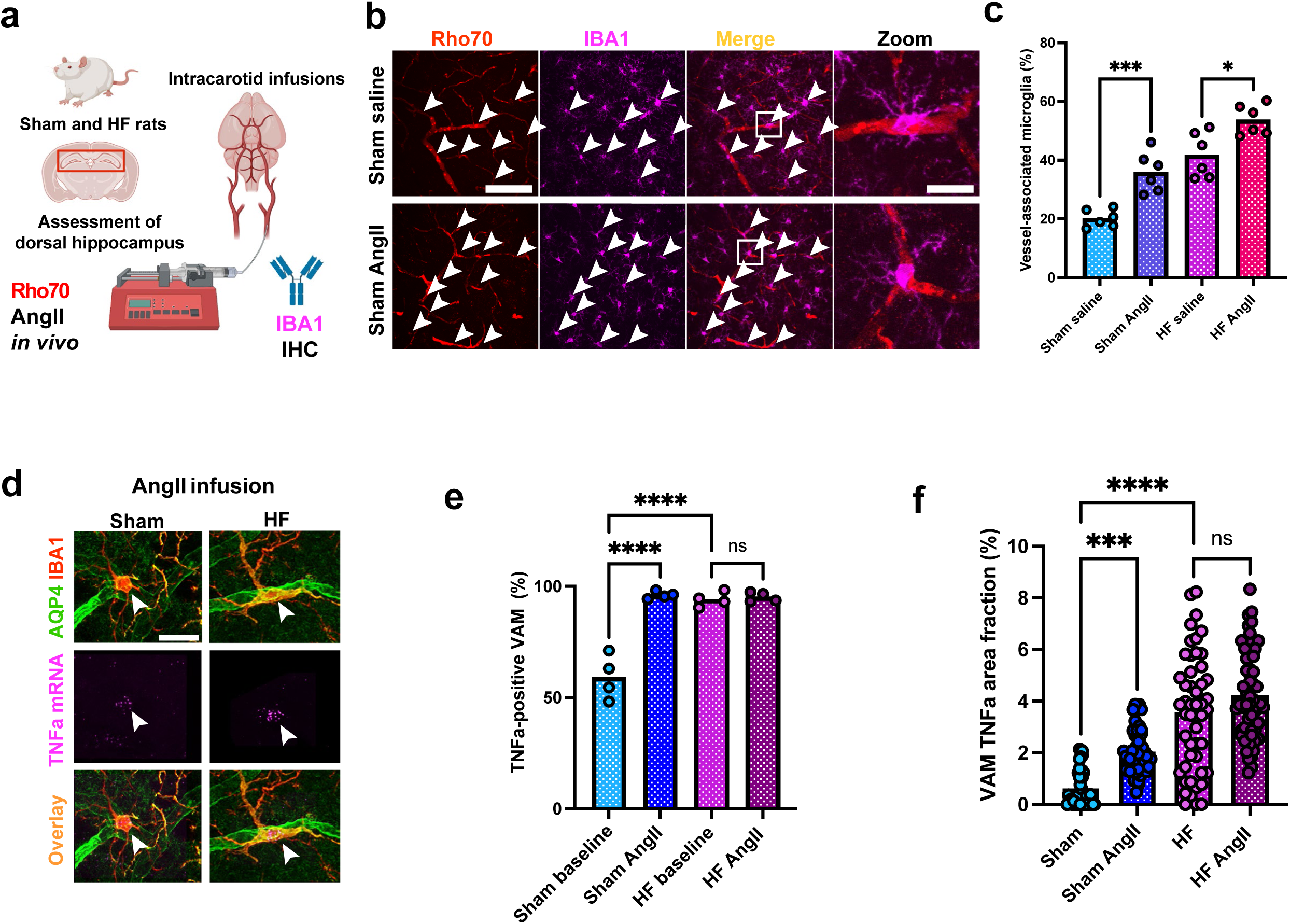
AngII-triggered and cytokine-mediated migration of microglia to blood vessels. **a** Schematic illustration of intracarotid infusions of Rho70 and AngII followed by immunohistochemical staining against IBA1 to label blood vessels and assess the numbers of parenchymal microglia and VAM. **b** Confocal images show parenchymal and VAM in a sham rat infused with saline and a sham rat infused with AngII; white arrowheads indicate the presence of VAM. High magnification show a VAM in both groups. Scale bars 100µm and 10µm. **c** Quantification of vessel-associated microglia after intracarotid AngII infusion (n=6 rats per group). **d** High magnification confocal images showing the co-localization of TNFα mRNA and vessel-associated, IBA1-positive microglia in sham and HF rats at baseline conditions and after intracarotid AngII infusion. White arrowheads highlight TNFα-positive, vessel-associated microglia. Scale bar 10µm. **e** Quantification of TNFα levels in individual vessel-associated microglia (single cell level area fraction in %) of sham and HF rats at baseline conditions after AngII infusion (N=53 cells from sham rats and N=57 cells from HF rats). **f** Quantification of TNFα-positive, vessel-associated microglia in sham and HF rats (n=4 rats per group) with and without AngII infusions.

TNFα is an important cytokine that stimulates microglia (and other immune cell) recruitment and migration ^77–79^. Thus, we performed a combination of RNAscope and immunohistochemistry to measure TNFα mRNA expression in VAM in sham and HF rats (**Figure 3d)** under this condition. We observed that the acute AngII infusion in sham rats increased both the proportion of VAM expressing TNFα (**Figure 3e**) as well as the levels of TNFα mRNA within individual VAM (**Figure 3f)**. In HF rats, we found that virtually all VAM were TNFα mRNA-positive under basal conditions (**Figure 3e)**, and while relative levels of TNFα mRNA were 2-fold higher in HF rats compared to sham (**Figure 3f),** AngII infusion failed to further increase expression in this group.

Finally, and to unequivocally demonstrate that AngII-AT1R signaling is necessary for the recruitment of microglia cells to the perivascular space in the context of HF, we performed AT1R blockage via administration of losartan, a commonly used AT1R antagonist delivered in the drinking water, as we recently reported ^19^ (**Figure 4a**). We found that the number of VAM in sham rats was not affected by losartan (**Figure 4b,c**). Conversely, losartan treatment significantly reduced the number of VAM by more than 2-fold in HF rats, normalizing their incidence to levels observed in sham rats (**Figure 4d-f**, HF: 45%, HF + Lo: 22%). Moreover, AT1R blockade in HF rats also normalized to the respective percentages of the different types of VAM (i.e., decreased in Type III and a concomitant increase in Type I (Chi-square test, p<0.0001, **Figure 4g**). Thus, these studies further support a key role for AngII-AT1R signaling in triggering VAM dynamics in the hippocampus of HF rats.

**Figure 4.**
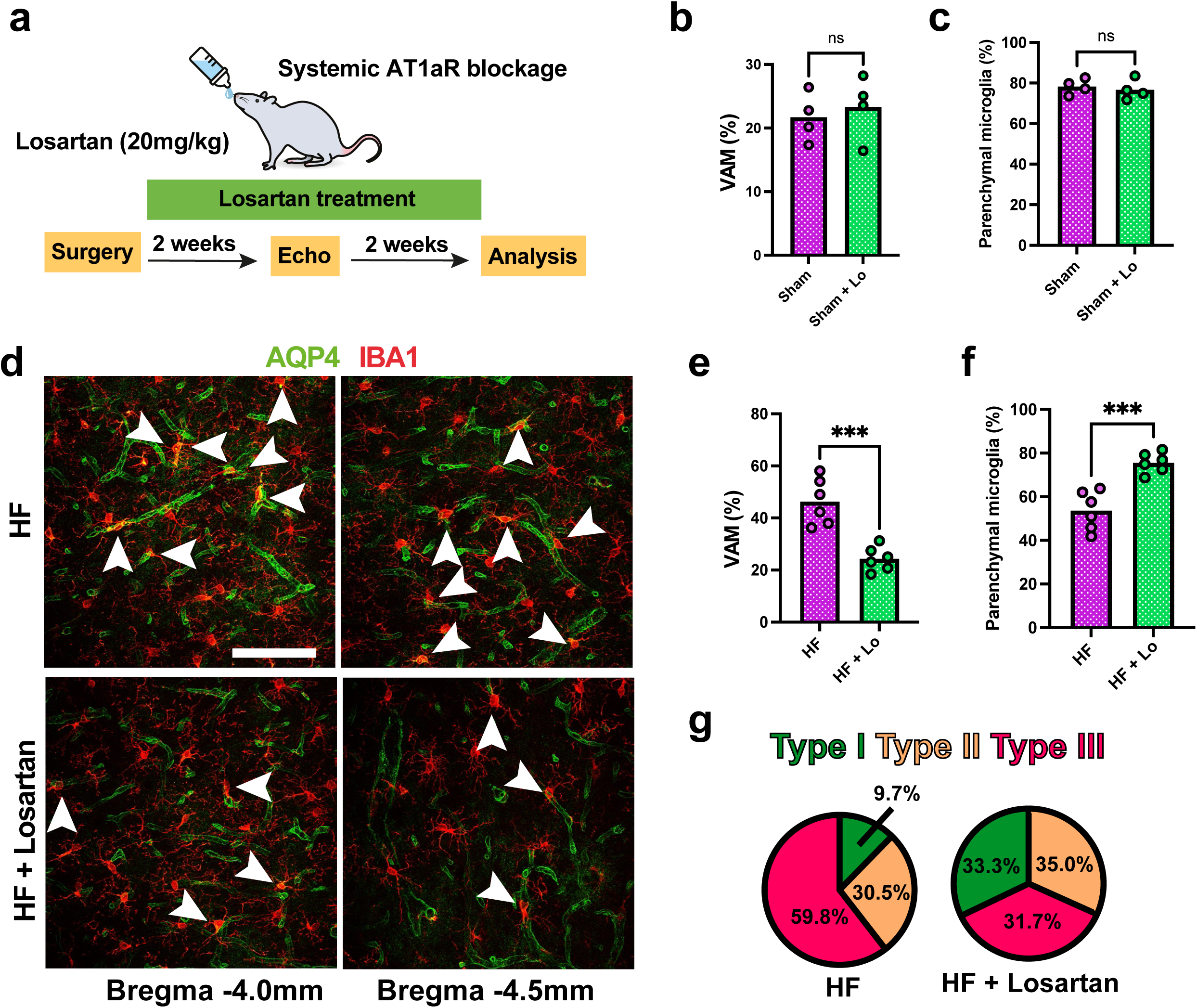
Losartan treatment blocks vessel migration in HF but not sham rats. **a** Schematic illustration of losartan administration via the drinking water in sham and HF rats. For two weeks after the surgery, losartan (20mg/kg) was supplied in the drinking water. Following echocardiographic assessment, rats were treated two more weeks before tissue collection and immunohistochemical staining and analysis. **b, c** Quantification of the number of vessel-associated and parenchymal microglia after losartan treatment in sham rats. **d** Confocal images show vessel-associated microglia with and without losartan treatment in HF rats at two different Bregma levels, scale bar 100µm. **e, f** Quantification of the number of vessel-associated and parenchymal microglia after losartan treatment in HF rats (n=6 rats per group). **g** Pie charts show the relative percentages of Type I-III vessel-associated microglia with and without losartan treatment in HF rats.

## Discussion

In this study, we used a combination of immunohistochemistry, three-dimensional reconstructions of the microglia-vascular interface, *in vivo* intracarotid infusion of vessel dyes and fluorescently-labeled AngII, RNAScope hybridization and pharmacological blockade of AngII-AT1aRs to study microglia-vessel dynamics in HF rats. Our main findings are: i) a 2-fold increase in vessel-associated microglia (VAM) in HF rats, a large proportion of which had processes than protruded into the vessel lumen; ii) a 2.5-fold increase in AT1aR expression in HF rats that was restricted to VAM, iii) recruitment of microglia to the perivascular space was induced by an acute systemic administration of AngII, and coincided with TNFα upregulation in VAM; and iv) administration of the AT1aR blocker losartan prevented the recruitment of microglia to the perivascular space, normalizing their levels and subtypes to those found in sham rats. These results highlight and support a highly dynamic microglial-vascular interface in the context of the pathophysiology of a highly prevalent cardiovascular disease.

### Microglial activation and neuroinflammation in heart failure: Role of the AngII-AT1aR signaling pathway

Microglia, traditionally considered as resident immune cells of the central nervous system, have been recognized for their versatile roles in various pathological conditions beyond the realm of neurological disorders ^25,26^. In the context of cardiovascular disease, and particularly heart failure (HF), a growing body of research has unveiled the significant involvement of microglia in orchestrating neuroinflammatory responses and contributing to the initial stages of the disease ^19,24,49,50^ and several studies have highlighted their pivotal contribution to the modulation of neuroinflammation and the early progression of cardiac dysfunction ^19,24,51,52^. Through the secretion of cytokines, chemokines, and reactive oxygen species, microglia are implicated in the amplification and propagation of neuroinflammatory signals during health and under various disease conditions ^25,53–56^. Thus, by influencing synaptic plasticity ^57–59^, neuronal excitability ^60,61^, and neurotransmission, microglia mediate neural circuitry modifications that can have profound effects on cardiovascular regulation and homeostasis. Attempting to harness these insights for therapeutic benefits, researchers have explored strategies targeting neuroinflammation to ameliorate HF symptoms ^19,23,49^. The rat ischemic HF model has been instrumental in evaluating the efficacy of potential interventions ^19,24,37,62–64^. Among the proposed interventions, microglia inhibitors have shown promise in attenuating neuroinflammatory responses, potentially dampening the progression of HF ^19,63,64^. AngII, a critical player in the pathophysiology of HF, occupies a central position in the cascade of events leading to neurohumoral activation ^30,52,65–68^ and cardiac dysfunction ^69–71^. As HF ensues, AngII levels surge, triggering a cascade of events that heighten sympathetic activity and prompt vascular constriction ^69–72^. This response is orchestrated to maintain blood pressure in the face of compromised cardiac ejection fraction and to initiate cardiac remodeling ^71,72^. Angiotensin receptor blockers such as losartan or angiotensin converting enzyme (ACE) inhibitors such as captopril are well-established pharmacological approaches for the treatment of patients with HF ^31,73,74^, and accumulating evidence supports that they can also act centrally to modulate microglia-mediated neuroinflammation ^19,38–40^. Our own previous work highlights the interaction between elevated circulating AngII levels and hippocampal microglial activation, neuronal dysfunction and apoptosis, and the contribution of this neuroinflammatory cascade to cognitive deficits in the ischemic rat HF model ^19^. We further showed that losartan reliably reversed all neuroinflammation-associated endpoints such as microglial morphology, astrogliosis, apoptosis and even cognitive impairment ^19^. Together, these results support a critical role for microglia in mediating AngII-induced neuroinflammation during HF.

### Vessel-associated microglia (VAM) dynamics in heart failure rats

Microglia-vascular interactions play a crucial role in maintaining the homeostasis and overall function of the brain ^25,26,41,48^ and the microglia-vascular interface has been the focus of numerous recent studies. Microglia-vascular interactions are multifaceted and are vital for the maintenance of the BBB that separates the circulating blood from the brain tissue ^41–43,45,75^. Microglia actively survey the brain microenvironment ^27,76^ and respond to changes in blood vessel integrity ^41,48^. In addition, they contribute to immune surveillance, modulate inflammation, and participate in the repair of damaged blood vessels ^41^. Importantly, recent work supports their ability to directly regulate local cerebral blood flow ^47,48^. Finally, dysregulation of microglia vessel interactions has been implicated in various neurological disorders, including neuroinflammatory and neurodegenerative diseases ^42,45^. Still, whether changes in microglia-vascular interactions also occur in the context of cardiovascular diseases, specifically HF,had yet to be determined.

Based on the rationale provided above, we focused on this work on microglial-vascular interactions in the dorsal hippocampus. We found that under normal physiological conditions, a small (20-30%) proportion of the hippocampal microglia is directly associated with the local microvasculature (i.e., vascular associated microglia, VAM), with either processes enwrapping the vascular wall, and/or their soma being in direct contact with the vessels (**Figure 2**). A similar proportion (20%) of VAM was previously reported in other studies ^41,48^.

An intriguing finding of our current study is the protrusion of the blood vessel wall by VAM filaments. This is in agreement with a recent work using electron microscopy that reported occasional microglia processes protruding through the vessel basal membrane and even obtruding through the vessel wall ^41^. Importantly, we found this phenomenon to be upregulated in HF rats (**Figure 2**). The precise role of vessel-protruding microglia remains elusive, but it seems plausible that VAM extend their filaments into the vessel structure as a means to “sense” circulating molecules including cytokines or even AngII. In fact, we found that more than 95% of VAM, particularly those with more extended physical contact with vessels (i.e. Type III) expressed AT1aR mRNA (**Figure 2**). These results suggest that hippocampal VAM could be positioned to sense the elevated levels of circulating AngII that occur during HF ^1,19,35,70^, thus being a key mechanism initiating a cascade of events leading to upregulation of cytokine expression and release (**Figure 2**), further microglial activation ^19^ and recruitment of additional microglia to the perivascular space (see model in **Figure 5**) in this disease state. This is supported by our study showing that an acute elevation of circulating levels of AngII was sufficient to increase the number of VAM in sham rats, as well as their expression levels of TNFα (**Figure 2**). In HF rats, the number of VAM and their TNFα expression levels were already higher compared to sham rats, but the AngII infusion was still able to further increase the number of VAM in this group. We previously showed that the AngII-AT1aR cascade stimulates the production of several cytokines in hippocampal microglia including C1q, IL1β and TNFα and that cytokine upregulation correlates with microglial deramification in HF rats ^19^. In this study, we focused on TFNα, given that it was previously shown to be an important signal that stimulates microglia (and other immune cell) recruitment and migration ^77–79^.

**Figure 5.**
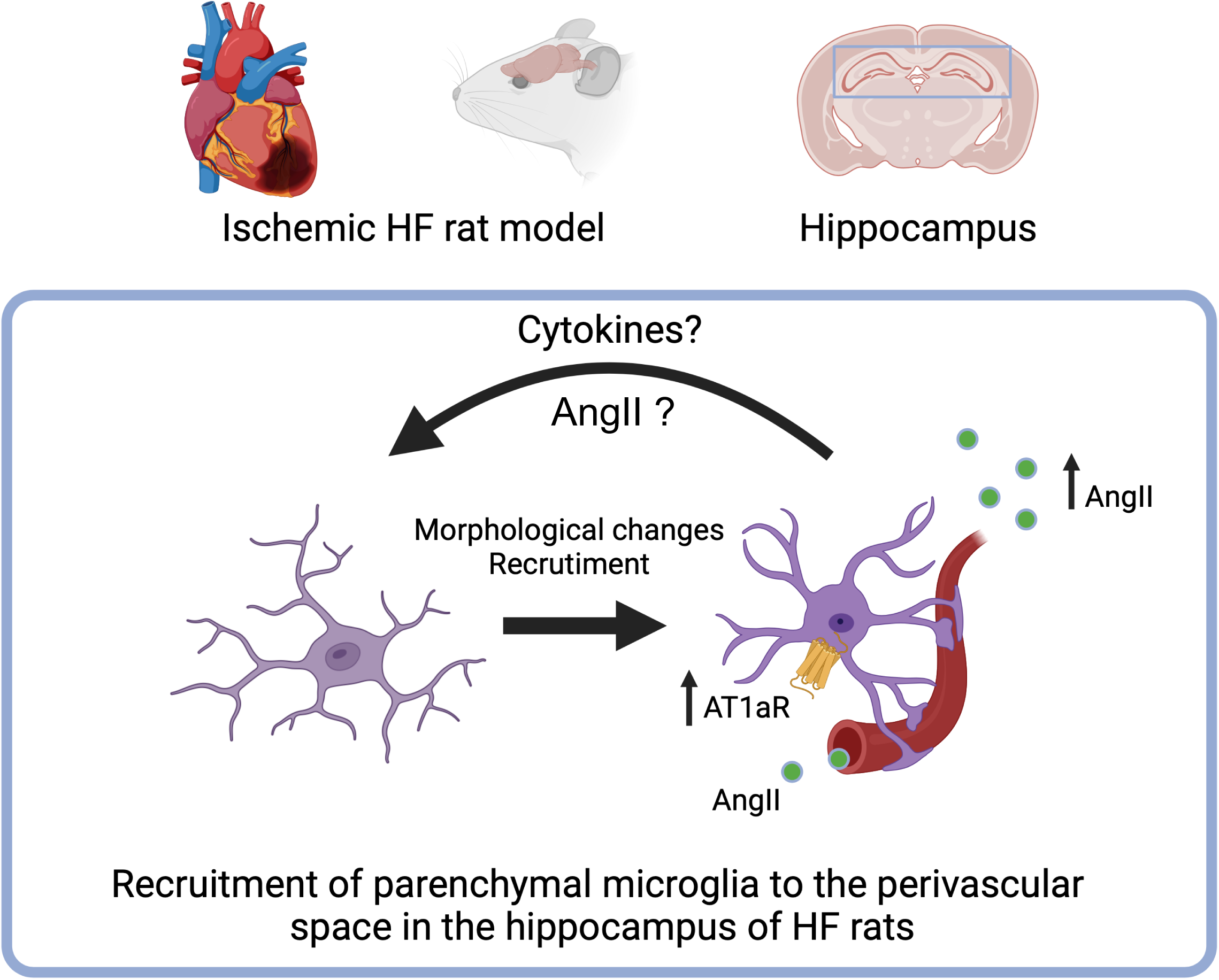
Working model for AngII-driven microglial recruitment during heart failure. Based on our current and previous findings, we hypothesize that elevated circulating AngII levels during HF and a subsequent microglia-specific AT1aR upregulation result in increased microglia-vessel interactions due to targeted and local microglial recruitment to the perivascular space. Upon activation, microglia produce and release, among other cytokines, TNFα, which further induces neuroinflammatory signaling, microglial activation and perivascular recruitment. This cascade might ultimately result in BBB deterioration and breakdown, allowing leakage of AngII into the hippocampal parenchyma, further exacerbating neuroinflammation and microglia perivascular recruitment.

Finally, and perhaps more importantly, we found that blocking the actions of endogenous AngII using a clinically relevant AT1aR blocker (losartan) following a clinically used route of delivery (oral administration) completely prevented the increase in hippocampal VAM in HF rats. Together, these results suggest that AngII-AT1aR signaling is critical for the recruitment of microglial cells to the vascular interface during HF.

The precise sequence of events leading to the AngII-AT1aR recruitment and activation of microglia during HF remains to be determined. One possible scenario, as summarized in the working model of **Figure 5**, is that the stable population of AT1aR expressing-VAM sense and react to the elevated circulating levels of AngII happening early in the onset of HF ^69,70^. This leads to their increased production of cytokines, including TNFα, overexpression of AT1aRs and a local neuroinflammatory response at the vascular interface, that ultimately results in AngII-mediated disruption of the BBB integrity, as we demonstrated in recent studies ^19,24^. We postulate that the combined action of BBB disruption with concomitant leakage of AngII into the hippocampal parenchyma, along with increased local production and diffusion of cytokines leads to further microglia activation and recruitment to the perivascular space, establishing thus a very disruptive positive feedback loop. This model is in line with our previous study, in which we demonstrated that microglial AngII signaling precedes cytokine production and that administration of losartan in HF rats significantly lowered various cytokine mRNA levels ^19^. Finally, our results showing that leakage of AngII through the disrupted BBB (a phenomenon also previously shown to be blocked by losartan^19^), was mostly bound to VAM (**Figure 2**) further supports the notion that leaked AngII also may play a role in the recruitment of parenchymal microglia to the perivascular space. Experiments done at early stages of HF are warranted to further confirm this hypothesis.

### Concluding remarks

Collectively, our findings in this study corroborate the key involvement of microglial AngII-AT1aR signaling in the pathophysiology of HF ^19^, while providing novel mechanistic insights contributing to this process. Specifically, our studies support the view that AT1aRs in VAM could be the site of initiation of the signaling cascade leading to AngII-mediated neuroinflammation, BBB disruption, hippocampal neuronal apoptosis, and cognitive deficits during HF ^19^. We propose that AngII-AT1aR signaling in VAM leads to further recruitment of parenchymal microglial to the microglial-vascular interface. Clearly, future *in vivo* studies are needed to determine whether physiologically relevant intravascular microglial protrusions exist, and whether microglia are able to directly monitor and sense circulating peptides such as AngII. Notwithstanding, our studies strongly support the notion that the microglial-vascular interface is highly dynamic, and that can be altered in the context of a prevalent cardiovascular disease. Thus, understanding the precise mechanisms and signals involved in this process is fundamental for the development of novel therapeutic targets for the treatment of cognitive and mood disorders associated to this disease.

## Material and methods

All experiments were approved and carried out in agreement to the Georgia State University Institutional Animal Care and Use Committee (IACUC) guidelines.

### Animals

We used male Wistar rats (5-7 weeks old at HF surgery, 180-200g, Envigo, Indianapolis, IN, USA) for all experiments (n=64). Rats were housed under constant temperature (22 ± 2°C) and humidity (55 ± 5%) on a 12-h light cycle (lights on: 08:00-20:00) *ad libitum* access to food and water.

### Heart failure surgery and Echocardiography

The ischemic HF surgical model and echocardiography was performed as previously described. The ejection fraction (EF) obtained via echocardiography was used as a key parameter to determine the degree of functional HF^10,24,80^. The mean EF values for sham and HF rats were 82.27±3.2 and 25.22±6.29, respectively (p<0.0001).

### Immunohistochemistry

Following pentobarbital-induced anesthesia (Euthasol, Virbac, ANADA #200-071, Fort Worth, TX, USA, Pentobarbital, 80mg/kgbw, i.p.), rats were first perfused at a speed of 20mL/min with 0.01M PBS (200mL, 4°C) through the left ventricle followed by 4% paraformaldehyde (PFA, in 0.3M PBS, 200mL, 4°C), while the right atrium was opened with an incision. Brains were post-fixed for 24 hours in 4% PFA at 4°C and transferred into a 30% sucrose solution (in 0.01M PBS) at 4°C for 3-4 days. For immunohistochemistry, 40 µm slices were cut using a Leica Cryostat (CM3050 S) and brain slices were kept in 0.01M PBS at 4°C until used for staining. Brain slices were blocked with 5% Normal Horse Serum in 0.01M PBS for 1h at room temperature. After a 15-min washing in 0.01M PBS, brain slices were incubated for 24h in 0.01M PBS, 0.1% Triton-X, 0.04% NaN_3_ containing different antibodies: 1:1000 of anti-IBA1 (polyclonal rabbit, Wako, 019-19741, Lot: CAK1997) or 1:500 of anti-AQP-4 (polyclonal rabbit, Alomone labs, AQP-004) at room temperature. Following 15-min washing in 0.01M PBS, sections were incubated in 0.01M PBS, 0.1% Triton-X, 0.04% NaN_3_ with 1:500 Alexa Fluor 488/594-conjugated donkey anti-rabbit (Jackson ImmunoResearch, 711-585-152, 705-585-147) for 4 hours at RT. Brain slices were washed again for 15 mins in 0.01M PBS and mounted using antifade mounting medium (Vectashield with DAPI, H-1200B/H-1500).

### RNAScope in situ hybridization

RNAScope reagents were bought from acdbio (PN320881). Nuclease-free water and PBS were acquired from Fisher Scientific. Brain tissues were handled according to the procedure outlined for Immunohistochemistry, utilizing nuclease-free PBS, water, PBS, and sucrose, in accordance with the instructions provided by the manufacturer. To evaluate the data, microglia were categorized as mRNA-positive if they exhibited three or more fluorescently-labeled voxel units within their individual cell bodies. Probes targeting AT1aR or TNFα mRNA (both C1, rat) were purchased from acdbio.

### Confocal microscopy and 3D analysis of microglia via Imaris

Confocal images were obtained using a Zeiss LSM 780 confocal microscope (1024×1024 pixel, 16-bit depth, pixel size 0.63-micron, zoom 0.7). Three-dimensional reconstruction of microglia was performed as previously described^24^. For the analysis of VAM and protruding microglia the surface-to-surface and surface-to-volume ratio feature of Imaris was used after prior enabling of object-to-objects statistics. Microglia were defined as VAM if they had overlapping surface with blood vessel of >2µm^2^ to avoid sampling of false positives as a result of reconstruction artifacts. Type I microglia displayed filamentous contacts, without somatic contact (i.e. distance from soma to vessel >1µm); Type II microglia displayed somatic contact with vessels (i.e. distance from soma to vessel <1µm) and Type III microglia displayed somatic contact with vessels (i.e. distance from soma to vessel <1µm) and extensive soma-vessel overlap defined as total surface-to-surface ratio > of >8µm^2^. Microglia were defined as vessel-protruding microglia when the overlapping volume (engulfed area) was >2µm^3^ to avoid sampling of false positives as a result of reconstruction artifacts. Whether the microglia had multiple vessel contacts or protrusions was not taken into account for the final analysis.

### Assessment microglial AngII_fluo_ uptake

Rats were anesthetized using a mixture of Ketamine and Xylazine (at concentrations of 60 mg/mL and 8 mg/mL respectively). A non-occluding catheter was inserted into the left internal carotid artery. Subsequently, AngII_fluo_ (at a concentration of 3µmol/L, obtained from Anaspec, CA) was infused at a rate of 2.86µl/g/rat and allowed to circulate for a duration of 30 minutes ^68^. After this period, the rats were decapitated, and their brains were fixed and sliced into sections that were 40µm thick using a Cryostat. These sections were prepared for confocal imaging. The software Imaris was utilized to identify and quantify the presence of microglial AngII_fluo_ using object-to-object statistics and quantification of engulfed AngII_fluo_ volume (in µm^3^). Microglia were considered to be AngII_fluo_ when they contained >2µm^3^ AngII_fluo_ to avoid sampling of false positives as a result of reconstruction artifacts.

### Losartan treatment

Rats with heart failure (HF) were assigned randomly to either the HF group or the HF + Losartan group. Losartan at a dose of 20mg/kg/day was administered in the drinking water, commencing one week after the HF surgery and continuing until the rats were sacrificed for analysis four to six weeks after the surgery.

### Statistical analyses

Statistical analyses were carried out using GraphPad Prism 9 (GraphPad Software, California, USA). To compare groups, various tests were employed, including Student’s t-test, Chi-square test, and one- or two-way analysis of variance (ANOVA), followed by Tukey post-hoc tests. Incidence of effects was compared using Chi-square tests. The results are denoted as mean ± standard error of the mean (SEM). Statistical significance was established at p < 0.05. In the corresponding figures, significance levels were indicated as * for p < 0.05, ** for p < 0.01, and *** for p < 0.0001.

## Data availability statement

The data that support the findings of this study are available from the corresponding author upon reasonable request.

## Acknowledgements

Schematic illustrations of Figures 2, 3 and 5 have been created using biorender.com.

